# Test-Retest Reliability of the Human Connectome: An OPM-MEG study

**DOI:** 10.1101/2022.12.21.521184

**Authors:** Lukas Rier, Sebastian Michelmann, Harrison Ritz, Vishal Shah, Ryan M. Hill, James Osborne, Cody Doyle, Niall Holmes, Richard Bowtell, Matthew J. Brookes, Kenneth A. Norman, Uri Hasson, Jonathan D. Cohen, Elena Boto

**Author notes:** **Correspondence to:** Dr Lukas Rier, Sir Peter Mansfield Imaging Centre, School of Physics and Astronomy, University of Nottingham, University Park, Nottingham NG7 2RD, United Kingdom.

## Abstract

Magnetoencephalography with optically pumped magnetometers (OPM-MEG) offers a new way to record electrophysiological brain function, with significant advantages over conventional MEG including adaptability to head shape/size, free movement during scanning, better spatial resolution, increased signal, and no reliance on cryogenics. However, OPM-MEG remains in its infancy, with significant questions to be answered regarding optimal system design and robustness. Here, we present an open-source dataset acquired using a newly constructed OPM-MEG system with a triaxial sensor design averaging 168 channels. Using OPM-optimised magnetic shielding and active background-field control, we measure the test-retest reliability of the human connectome. We employ amplitude envelope correlation to measure whole-brain functional connectivity in 10 individuals whilst they watch a 600 s move clip. Our results show high repeatability between experimental runs at the group level, with a correlation coefficient of 0.81 in the theta, 0.93 in alpha and 0.94 in beta frequency ranges. At the individual subject level, we found marked differences between individuals, but high within-subject robustness (correlations of 0.56 ± 0.25, 0.72 ± 0.15 and 0.78 ± 0.13 in theta, alpha and beta respectively). These results compare well to previously reported findings using conventional MEG; they show that OPM-MEG is a viable way to characterise whole brain connectivity and add significant weight to a growing argument that OPMs can overtake cryogenic sensors as the fundamental building block of MEG systems.

## INTRODUCTION

Magnetoencephalography using optically-pumped magnetometers (OPM-MEG) is an emerging technique to image human brain function (see Brookes et al. (2022) for a review). As with conventional MEG, electrophysiological activity is assessed non-invasively by measuring magnetic fields at the scalp surface generated by neural currents (Cohen, 1968). However, unlike conventional MEG which employs arrays of cryogenically cooled sensors (Cohen, 1972; Hamalainen et al., 1993), OPM-MEG uses small and lightweight detectors – OPMs – which do not require cooling. Cryogenic temperatures place significant restrictions on MEG system design, requiring large and cumbersome instrumentation, with sensor arrays fixed rigidly inside a one-size-fits-all helmet. OPMs lift these restrictions leading to several advantages, for example, sensors can be positioned closer to the head, increasing signal amplitude and spatial resolution; lightweight sensors can be mounted in a wearable helmet, ostensibly enabling free subject movement during data acquisition; freedom to place sensors anywhere means OPM-MEG can, in principle, adapt to head size, enabling lifespan compliance; and systems are relatively simple to build, site, and operate.

The capability of OPMs to measure the MEG signal has been shown extensively, e.g. (Boto et al., 2017; Johnson et al., 2010; Kamada et al., 2015; Sander et al., 2012; Xia et al., 2006), and OPM arrays have been developed which can image brain function accurately and, in some cases, with whole head coverage (Feys et al., 2022; Hill et al., 2020; Iivanainen et al., 2019; Johnson et al., 2013; Nardelli et al., 2020; Seymour et al., 2021). Improved data quality has been shown in both theory (Boto et al., 2016; Iivanainen et al., 2017) and practice (Boto et al., 2017), including during subject movement (e.g. Boto et al. (2018)), though recording during active movement critically depends on background field control (Holmes et al., 2018, 2020; Rea et al., 2021). Applications in children are also beginning to emerge (Hill et al., 2019) with exciting clinical possibilities (Feys et al., 2022). In sum, emerging OPM-MEG systems offer new opportunities which are not possible using conventional neuroimaging. However, OPM-MEG remains nascent technology – there are only a small number of systems worldwide and few have been tested for robustness. The best ways to design OPMs, sensor arrays, and magnetic shielding (which controls background fields) are not yet settled, and there is relatively little open-source data available from OPM-MEG systems. In this paper, we aimed to evaluate a recently developed triaxial OPM-MEG instrument (Boto et al., 2022; Rea et al., 2022) via quantitative assessment of test-retest reliability for the measurement of human connectomics. We further intended to generate an open-source dataset to allow other researchers to assess OPM-MEG capabilities.

Our system employs triaxial OPMs which allow independent measurement of the magnetic field along three orthogonal axes (Shah et al., 2020). Despite a slightly higher noise floor compared to conventional (single or dual axis) OPMs, triaxial sensors are an effective means to interrogate the MEG signal (Boto et al., 2022). They also allow increased total signal strength (i.e. each sensor makes three measurements of field)(Brookes et al., 2021; Rea et al., 2022), improved ability to differentiate brain activity from background fields (Brookes et al., 2021; Rea et al., 2022; Tierney et al., 2022), more uniform coverage in infants (Boto et al., 2022) and optimised calibration. In addition to the triaxial array, our system includes magnetic shielding which operates in active feedback configuration (Rea et al., 2021). This means that both low-frequency drifts in the background field and the static (i.e. time-invariant) magnetic field inside a magnetically shielded enclosure (MSE) are suppressed (Holmes et al., 2020), so data are collected in close to “zero” field. A prototype (90-channel) version of this system, with limited coverage of the head, has been previously demonstrated (Rea et al., 2022).

Over the last two decades, functional connectivity has emerged as an important means to characterise brain health. Data from functional magnetic resonance imaging (fMRI) and MEG have shown that even with the brain “at rest”, spatially separate but functionally related regions communicate to form networks. Some networks are associated with sensory processes, others with higher-level functions like attention or cognition. These networks are key to healthy brain function and are often perturbed in neurological and psychiatric disorders. MEG offers multiple means to measure connectivity (O’Neill et al., 2015a) and provides a tool to better understand the neural substrates that underlie communication (Sadaghiani et al., 2022). In addition, the exquisite time resolution of MEG allows us to look for dynamic changes in network connectivity, at the scale of seconds (O’Neill et al., 2015b) and milliseconds (Baker et al., 2014). Consequently, the accurate and reliable measurement of network connectivity plays a critical role for any MEG system. However, connectivity measurement is also a challenge: the distributed nature of networks requires whole-head coverage and since mathematical techniques to characterise connectivity (particularly in the resting state) must be applied to unaveraged data, high-fidelity recordings are paramount.

Functional connectivity has been measured previously using OPM-MEG, during tasks and in the resting state (Boto et al., 2021), with results comparable to a conventional MEG system. However, this was with an early whole-head instrument (50 radial channels) and test-retest robustness was not assessed. Even with conventional MEG, the test-retest reliability of connectivity is challenging, for example, Colclough et al. (2016) showed that whilst at the group level (∼30 subjects) repeatability of connectome estimation is excellent (>95%, based on amplitude envelope correlation), at the individual level reproducibility is closer to 60% (within-subject), and this drops further (<50%) for between-subject comparisons. Liuzzi et al. (2017) showed within-subject test-retest correlations of just ∼58% using conventional MEG, but also that longer MEG recordings (10 mins relative to 5 mins) and immobilising the head to prevent movement relative to the sensor array significantly improved consistency, to >70%. The extent to which remaining differences between runs are due to instrumentation, or real differences in brain activity, is unknown. The extension of such metrics to OPM-MEG would be a significant step forward.

In this paper, we characterise the robustness of connectomics using OPM-MEG. To maximise the chances of high reliability we used 10-minute recordings and, to provide a degree of consistency in brain activity, participants watched a movie clip during the scan. We chose a movie-viewing paradigm that has been used previously to assess relationships between fMRI, EEG and electrocorticography (ECoG) (Haufe et al., 2018). This standard task not only facilitates our objective to measure robustness but also provides a new open-source resource with direct equivalence to existing data (Haufe et al., 2018). We quantitatively assess consistency between separate experimental runs and thus provide a benchmark for the reliability of connectivity measurement using OPM-MEG.

## MATERIALS AND METHODS

### Subjects and Experimental Paradigms

10 participants (4 identified as female; 6 as male, all right-handed) gave written informed consent to participate in the experiment, which was approved by the University of Nottingham Medical School Research Ethics Committee. Each participant was scanned twice. During both recordings, participants watched a 600 s clip of the movie “Dog Day Afternoon” (Haufe et al., 2018; Honey et al., 2012; Lumet, 1975). Subjects remained seated and continued wearing the sensor helmet between scans (so that a single co-registration of sensor geometry to brain anatomy could be used for both measurements, reducing co-registration error). Before the MEG recording, a field-mapping and nulling procedure (Rea et al., 2021) was carried out to control the background magnetic field (see below). MRI scans (acquired using a Phillips Ingenia 3 T MRI system running an MPRAGE sequence, with 1-mm isotropic resolution) were also acquired for all ten participants.

### The OPM-MEG system

We used an OPM-MEG system averaging 168 channels, constructed from triaxial OPMs, each yielding three independent channels per sensor (Boto et al., 2022; Shah et al., 2020) (QuSpin, Inc. Colorado, USA). The sensors were spaced evenly around the scalp and mounted in a 3D-printed lightweight helmet (Cerca Magnetics Ltd., Nottingham, UK), affording approximately whole-head coverage. The helmets came in multiple sizes and the best-fitting helmet was chosen for each participant. Outputs of all channels were recorded via a digital data acquisition (DAQ) system (National Instruments, Austin, TX, USA). Participants were seated on a patient support inside an OPM-optimised magnetically shielded room (MSR) (Cerca Magnetics Ltd., Nottingham, UK). The MSR comprised four layers of mu-metal and one layer of copper, and was equipped with degaussing coils (Altarev et al., 2014) to reduce the magnetisation of the mu-metal layers – the static background field at the centre of this room following degaussing of the inner-most layer is typically ∼3 nT. To further control the field, an array of four (QuSpin, first generation) reference OPMs was placed immediately behind the subject to measure background field fluctuations, and a set of biplanar electromagnetic coils were placed on either side of the participant, which enabled the generation of all three uniform fields and 5 independent linear gradients in a 40 × 40 × 40 cm^3^ region enclosing the subjects head. The room also housed a motion tracking system comprising 6 cameras (OptiTrack Flex 13, NaturalPoint Inc., Corvallis, OR, USA) placed around the MSR, which recorded the movement of infrared-retroreflective markers attached to the bi-planar coils (as a static reference) and the sensor helmet (to monitor head movement). The OPM sensors, DAQ, storage, and background field compensation were controlled via a single (“acquisition”) PC. A second (“stimulus”) PC controlled the movie and motion tracking. The visual display was achieved via projection through a waveguide onto a back-projection screen. We used a View Sonic PX748-4K projector positioned outside the MSR, and the screen was placed ∼80 cm in front of the subject. The movie was presented at a visual angle of ∼13 degrees horizontally and ∼9 degrees vertically. Audio was presented through a set of speakers mounted outside the MSR and connected to a waveguide via a plastic tube. A schematic of the full system is shown in Figure 1A; a photograph of a participant wearing the system is shown in Figure 1B.

**Figure 1:**
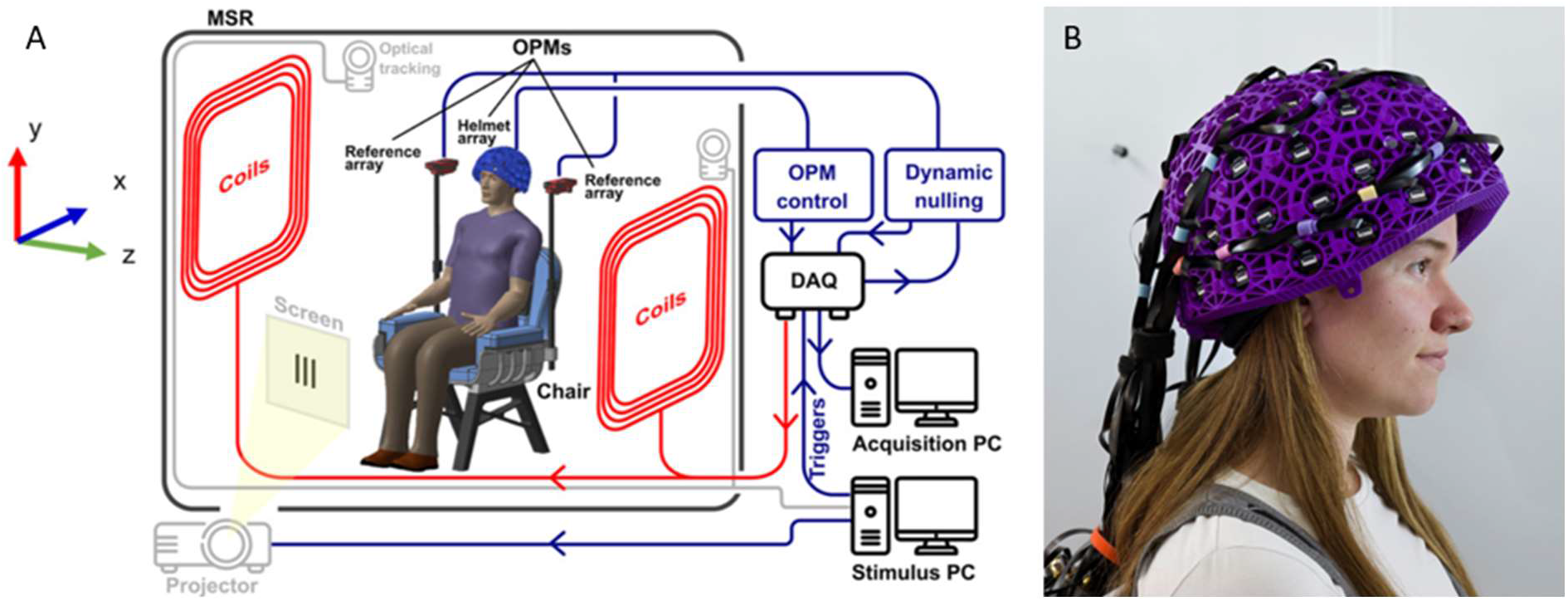
the OPM-MEG system. A) Schematic adapted from Rea et al. (2022b) showing the OPM-MEG system setup. B) A photograph of a participant wearing the OPM-MEG system.

### Magnetic field control

We used the field-control techniques originally described by Rea et al. (Rea et al., 2021). Briefly, after positioning participants in the MSR, the MSR door was closed and the inner mu-metal layer was degaussed. The reference OPM array was used to sample background field fluctuations, and the data were fed back to the (calibrated) bi-planar coils which generated an equal and opposite field. In this way, slow (<3 Hz) changes in the 3 uniform components of the field and the three gradients varying in *z* (from the participant’s right to left), were stabilised (Holmes et al., 2020). This left only a static (i.e. temporally invariant) background field which was measured via a nulling procedure in which participants executed translations and rotations of their heads. The motion of the helmet was tracked for 60 s and 5 OPMs (15 channels) were used to sample the changes in magnetic field induced by the movement. These data were combined, and the background field was modelled using spherical harmonics. The calculated three homogeneous field components and 5 linear gradients were then compensated using the bi-planar coils. This nulling process was repeated twice (to iteratively improve the estimate) and the modelling was repeated a third time to estimate the magnitude of the background field in which the experimental MEG data were captured.

### Data collection and co-registration

A total of 600 s of OPM data were recorded for each participant and each run of the experiment. All OPM channels were sampled at 16-bit resolution with a sampling rate of 1200 Hz. At least once per scanning day, a 90-s measurement with no subject present – termed ‘empty-room noise data’ – was also acquired to verify that the system was working appropriately. This meant that in total, seven empty room recordings were also available for analysis (see below) alongside the MEG data.

Immediately following MEG data acquisition, two 3D digitisations of the participants’ heads were acquired using an optical imaging system (Einscan H, SHINING 3D, Hangzhou, China) – the first with the helmet on and a second with the helmet removed and a swimming cap used to flatten any hair. A 3D surface representing the face and scalp was also extracted from the anatomical MRI. These data were used to enable co-registration of the MEG sensor geometry to brain anatomy. Briefly, the two optical digitisations were segmented, leaving only points around the face which were then aligned. The second optical digitisation (with the helmet removed) was then aligned to the surface extracted from the MRI. These two steps enabled knowledge of the helmet position relative to the brain. The locations and orientations of the OPMs, relative to the helmet, were known from the 3D printing process and the addition of this information enabled complete co-registration (see also Zetter et al. (Zetter et al., 2019) and Hill et al. (Hill et al., 2020)). This co-registration was used subsequently to facilitate forward modelling of the magnetic fields generated by current dipoles in the brain.

## Data Analysis

### Pre-processing and artefact correction

OPM-MEG data for each experiment (and the corresponding empty noise recordings) were notch-filtered at the mains frequency (50 Hz) using a second-order infinite impulse response filter (Q-factor of 35 at -3dB), and band-pass filtered (1-150 Hz) using a 4^th^-order, zero-phase-shift Butterworth filter. The filtered data were inspected visually for noisy and/or failed channels which were removed. On average, 152±3 clean channels were included in the final analyses. Each experimental recording was divided into 5-s epochs which were characterised as ‘good’ or ‘bad’: epochs were inspected visually and trials containing visible motion or muscle artefacts were marked as bad. Additionally, an automatic thresholding procedure was used to remove trials containing large artefacts: specifically, the standard deviation of the 1-150 Hz data within each epoch was calculated independently for each channel. Epochs containing more than one channel with a standard deviation exceeding 3 standard deviations from the mean (calculated over all time) were marked as ‘bad’. On average, 17±5 bad trials (18±4 in run 1, 17±5 in run 2) were removed resulting in 513±24 s of clean data (mean ±std. deviation across recordings). Independent component analysis (ICA) (FieldTrip implementation – (Oostenveld et al., 2011)) was used to identify and remove ocular and cardiac artefacts: the data were decomposed into a number of components equal to the channel count and visual inspection of component time-courses used to identify the artefacts. Finally, homogeneous field correction (HFC) (Tierney et al., 2021) was applied to attenuate interference from distal sources of magnetic field (which manifest as approximately uniform over the OPM array).

### Source reconstruction

A beamformer (Robinson and Vrba, 1999) was used for source reconstruction. The brain was parcellated into 78 cortical regions, defined by the Automated Anatomical Labelling (AAL) atlas (Gong et al., 2009; Hillebrand et al., 2016; Tzourio-Mazoyer et al., 2002). This was achieved by co-registering the AAL atlas to individual brain space using FLIRT in FSL (Jenkinson et al., 2002; Jenkinson and Smith, 2001). The coordinates of the centre of mass of each AAL region were determined and forward fields for each resulting location were calculated. The forward calculation was implemented using a dipole approximation and a single shell volume conductor model based on a head shape extracted from the anatomical MRI using FieldTrip (Nolte, 2003). Source reconstruction was repeated using data covariance based on broad-band data (1-150 Hz) and six bands of interest (BoIs) encompassing the canonical theta (*θ*: 4-8 Hz), alpha (*α*: 8-12 Hz) and beta band (β: 13-30 Hz), as well as three ranges within the gamma band (*γ*_1_: 30-40 Hz, *γ*_2_: 35-45 Hz, *γ*_3_: 40-48 Hz). Pre-processed data were band-pass filtered to each BoI using a 4^th^-order, zero-phase-shift Butterworth filter and covariance matrices constructed using data recorded throughout the whole experiment. Covariance matrices were regularized using the Tikhonov method by adding 5% of the maximum singular value of the unregularized matrix to the leading diagonal. The forward fields and data covariance were used to calculate beamformer weighting parameters, where source orientation was determined as the direction of maximum beamformer projected signal amplitude (Sekihara et al., 2004). Multiplication of the weighting parameters with the data resulted in 7 electrophysiological time series (one for each frequency band) at each of the 78 regions defined by the AAL atlas. This was repeated for every subject and independently for each experimental run.

### Spectral Power

To visualise the spectral content across AAL regions, and to examine the consistency of the beamformer projected signals between the two experimental recordings, we performed two analyses. First, we took the broadband (1-150 Hz) beamformer projected data, normalised by its standard deviation, and filtered to each BoI (using a 4^th^-order, zero-phase-shift Butterworth filter). The variance of the filtered data thus offered an estimate of the relative contribution of each BoI to the signal in a specific region. Applying this to all BoIs and regions allowed us to construct maps showing the spatial signature of the relative contribution of each band to the total signal for each AAL region. Secondly, for each region, we took the broadband beamformer projected data and used Welch’s method to estimate the power spectral density (PSD). We also applied the same beamformer weights to project the empty room noise data. This enabled visualisation of not only the consistency of the PSD across recordings but also of the relative contribution of empty room noise. We estimated the fractional difference in spectral power between runs as the square root of the sum of squared differences between PSDs, for runs one and two, normalised by the total integral of the overall mean PSD.

### Functional Connectivity

Functional connectivity between all pairs of AAL regions and for each BoI was calculated using amplitude envelope correlation (AEC) (Brookes et al., 2011; O’Neill et al., 2015a). The narrow-band beamformer projected data were taken for two regions, and pairwise orthogonalisation was applied to reduce the effect of source leakage (which is known to affect estimates of functional connectivity (Brookes et al., 2012; Hipp et al., 2012). Following orthogonalisation, a Hilbert transform was applied to the data from each region and the analytic signals were calculated. The absolute value of the analytic signals was then used to determine the “Hilbert Envelope” (i.e. the instantaneous amplitude envelope of band-limited oscillations for each of the two regions). These envelopes were down-sampled temporally from 1200 to 120 Hz and the Pearson correlation coefficient between the envelopes was used to quantify functional connectivity. This procedure was applied to all (78^2^ – 78 =) 3003 possible region pairs within the AAL parcellation, resulting in a whole-brain connectome. The analysis was run independently for each experimental run, participant and BoI.

To visualise the connectome matrices, they were normalised by dividing each matrix element by the square root of the mean of all squared matrix elements and averaged across subjects (preventing a single subject with high connectivity values from dominating the group average). This produced a group mean connectome for the first and second experimental runs, and each BoI, separately. The matrices were plotted, and in addition, thresholded to keep only the 150 strongest connections which were plotted as lines within a glass brain. We also assessed average global connectivity – the mean across matrix elements, before normalisation – and the mean paired difference in global connectivity between runs

We quantified the reliability of the group-average connectomes by calculating the Pearson correlation coefficient (using only matrix elements above the leading diagonal) between the subject averages for the two runs (separately for each BoI). We also assessed the influence of group size; for sample sizes of *N* = {2, 3, … 9}, all possible combinations of subjects were drawn, and average connectomes calculated. We then measured the between-run correlation. By plotting the mean and standard deviation of these correlations for each *N*, we were able to estimate the trajectory of between-run consistency with increasing N.

### Inter-individual differences

In addition to group analyses, we examined connectivity at the individual level and the sensitivity of our OPM-MEG system to differences between participants. With 10 subjects, each scanned twice, there are 100 independent comparisons between run 1 and run 2 that can be made at the individual level; 10 within-subject comparisons and 90 between-subject comparisons. For every possible comparison, we measured the Pearson correlation between vectorised matrices (again using only elements above the leading diagonal). We analysed these in two ways. First, we averaged the within and between subject correlations, computed the difference in the mean, and tested to see if this difference was significant using a Monte-Carlo test. Specifically, we randomly switched which 10 values were chosen as the within-subject correlations; doing this for 100,000 iterations enabled the construction of an empirical null distribution and allowed us to estimate whether the real difference could have occurred by chance. Second, we performed a “neural fingerprinting” analysis. For every subject, there is one within-subject comparison and 9 between-subject comparisons – one might expect that the correlation coefficient for the within-subject comparison should be higher than the other nine values. If it is, that subject can be said to be successfully identified. By repeating this ten times, we were able to assess how many (out of ten) subjects could be correctly identified based on their MEG connectome data.

The data presented here have been made publicly available (Rier et al., 2022), enabling free access to OPM-MEG data for the neuroimaging community – a core aim of the current study.

## RESULTS

Our field modelling showed that – following degaussing of the MSR walls and the application of average coil currents – the magnitude of the uniform magnetic field components inside the MSR was 0.54 ± 0.33 nT, with linear gradients of 1.70 ± 0.75 nT/m. These values dropped to 0.19 ± 0.17 nT and 0.63 ± 0.69 nT/m for the second field mapping. Comparable conditions were achieved previously (Rea et al., 2021).

### Power spectral density

Figure 2A shows the spatial signature of spectral power in different bands of interest during the task. As expected, alpha oscillations dominate the signal in occipital areas, with similarly high contributions stretching forward to the parietal lobes. Beta oscillations were highest in sensorimotor regions. Theta oscillations were approximately uniform across the whole head whilst gamma oscillations were most prominent in the frontal areas.

**Figure 2:**
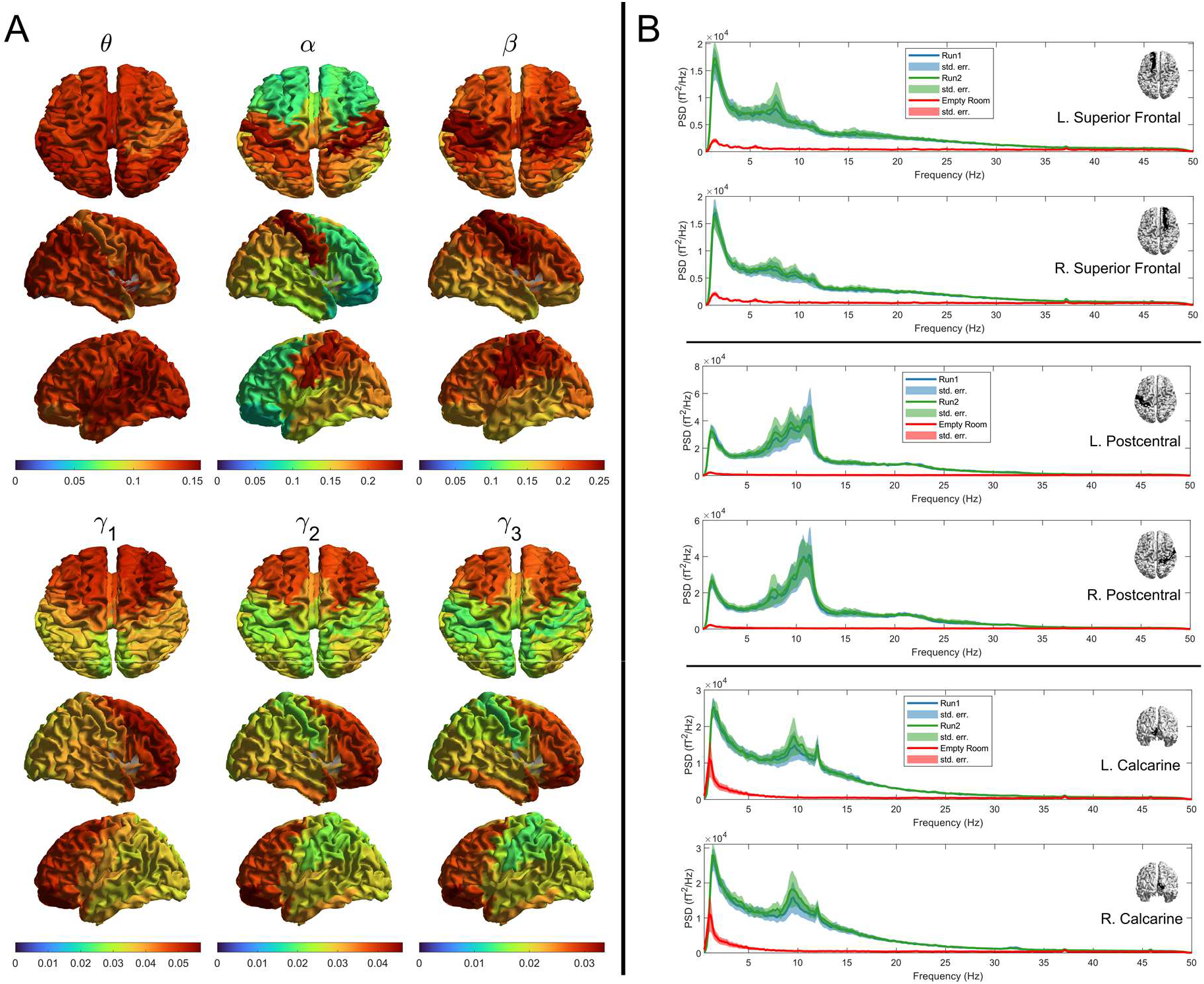
Spectral Power. A) Brain plots showing the mean spatial topographies of relative spectral power averaged across subjects and runs in theta (*θ*: 4-8 Hz), alpha (*α*: 8-12 Hz), beta (β: 13-10 Hz) and overlapping sub-bands of the gamma band (*γ*_1_: 30-40 Hz, *γ*_2_: 35-45 Hz, *γ*_3_: 40-50 Hz). B) Broad-band power spectra for example regions indicated in the corresponding insets. Blue and green lines represent the group average spectra for the first and second runs respectively. Shaded areas correspond to the standard error across subjects in each run. Red lines indicate the average spectrum obtained by projecting data recorded in the empty MSR using the beamformer weights estimated for each subject and run at each of the chosen locations.

Figure 2B shows example power spectral density plots for six selected AAL regions – left and right superior frontal, postcentral and calcarine cortex. In all cases, PSD for run 1 is shown in blue, run 2 in green, and red shows the PSD of the beamformer-projected empty room noise. In agreement with Figure 2A there are differences between regions – for example, elevated beta power is observed in the sensorimotor regions and prominent alpha peaks exist in the occipital areas. Most importantly, note the high level of consistency between experimental runs. The square root of the summed square of the differences between the PSD values for the two runs over the integral of the average of the two was found to be 4±1% (mean ± std. deviation) when averaged over the 78 AAL regions; when examining the variation of this difference across brain regions, it was dominated by differences in occipital, parietal and temporal lobes. The largest difference between runs was elevated alpha power in run 2, compared to run 1 (Wilcoxon sign rank test, p=0.0039). Differences in power in the other bands did not survive multiple comparisons correction.

For frequencies below around 60 Hz, the projected empty room noise was lower than the signal, implying a good ratio of signal to sensor noise/interference. On average, the ratio of signal to noise (i.e. the ratio of the green/blue lines to the red line) was 14±8 for θ, 24±18 for α, and 8±4 for β. However, this decreased to 2.7±0.9 for γ_1_, 2.0±0.5 for γ_2_ and 1.8±0.4 for γ_3_, demonstrating how the signal amplitude approaches the sensor noise level with increasing frequency. This is an important point for OPM-MEG sensor design and will be addressed later in the discussion.

### Functional connectivity at the group level

Figure 3A shows group-level connectome results. Connectome matrices are shown alongside glass brain plots in which the lines show the spatial signature of the strongest 150 connections, whilst the blue circles show connectivity strength (i.e. the sum of the connectome matrix in one direction, representing how connected that brain region is to all other regions). Results for all BoIs are shown. As expected, the spatial signature of connectivity is different in different frequency bands: The alpha band is dominated by occipital, temporal and posterior parietal connections; the beta band has the highest connectivity strength in sensorimotor regions, with additional frontoparietal and occipital projections. The low gamma band also highlights a strong sensorimotor network. The theta band has strong posterior connections but with some frontal projections, whilst the two highest frequency (gamma) bands appear to identify frontal and superior parietal connections. These spatial signatures are in good agreement with those found using conventional MEG (Hunt et al., 2016). Figure 3B shows the mean global connectivity (averaged over the whole connectome matrix) for the two runs, for each band. Figure 3C shows the difference between runs (i.e. a paired subtraction of global connectivity within each subject, averaged across subjects). In all cases, the bar heights show the mean value and error bars show the standard deviation across participants.

**Figure 3.**
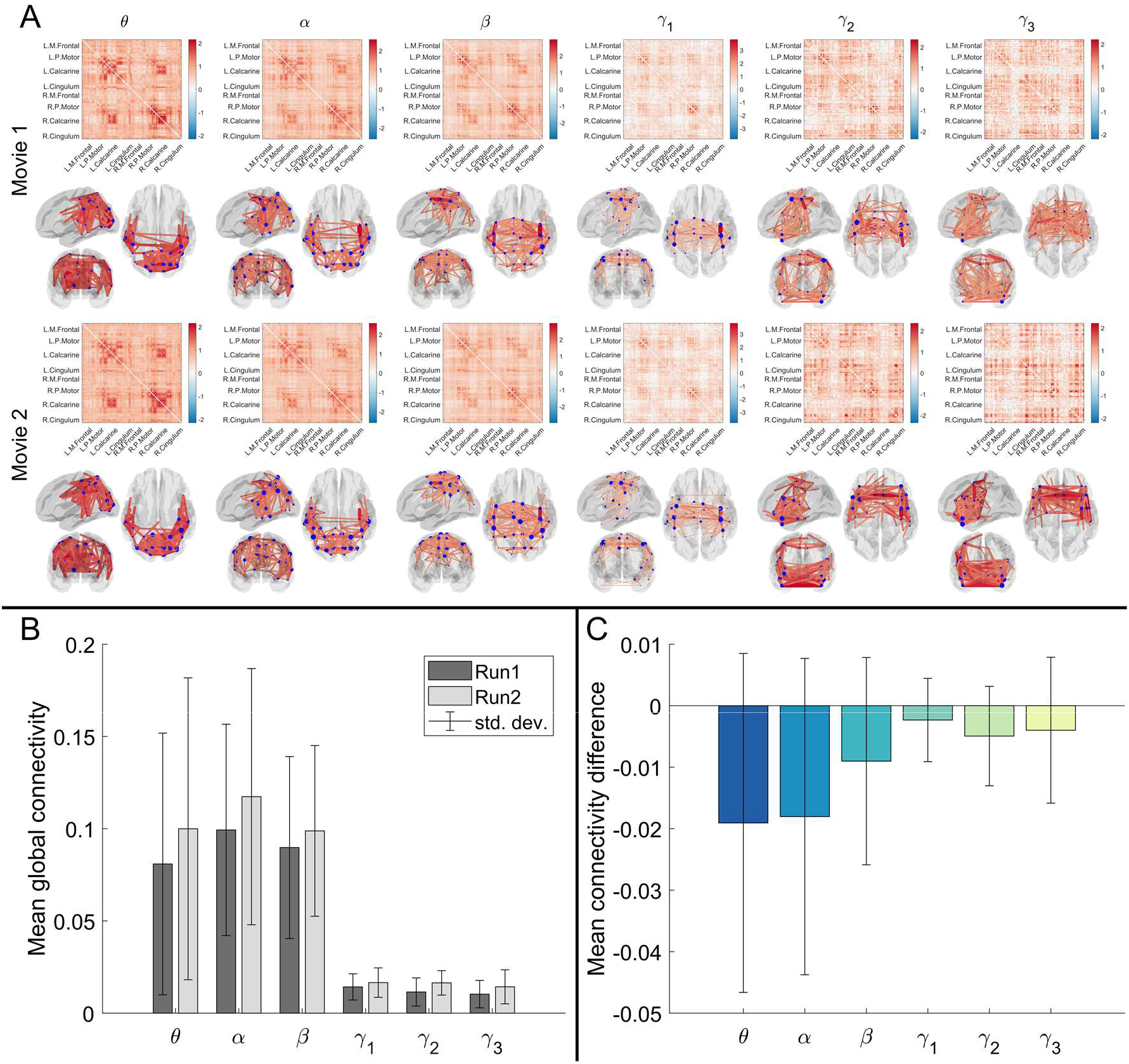
Group average amplitude envelope correlation and between-run consistency. A) Group average connectomes calculated using amplitude envelope correlation for 2 separate experimental runs, in theta (θ: 4-8 Hz), alpha (α: 8-12 Hz), beta (β: 13-10 Hz) and overlapping sub-bands of the gamma band (γ_1_: 30-40 Hz, γ_2_: 35-45 Hz, γ_3_: 40-50 Hz). Glass brains show the strongest 150 connections B) Mean global connectivity across subjects for each run and frequency band. C) Mean connectivity difference between runs for each frequency band.

There is a slight trend towards higher global connectivity in the second experimental run compared to the first, though this did not reach significance (a paired Wilcoxon sign rank test on the difference values suggested p-values of 0.05, 0.06 and 0.23 for theta, alpha and beta bands respectively – no measures survived a multiple comparison correction across bands). Most importantly, in the theta, alpha, beta and low gamma bands there is marked similarity in the structure of the connectome matrix across the two separate experimental runs.

This is formalised in Figure 4, where panel A shows all matrix elements from run 1 plotted against all matrix elements for run two. Between-run correlation coefficients are shown in panel B as a function of frequency band. Consistency between runs peaks in the beta band with a correlation coefficient of 0.935. Correlation is also high for alpha (0.929) and theta (0.814) but declines with increasing frequency to 0.714, 0.599 and 0.54 for the three gamma bands. Figure 4C shows the relationship between sample size (i.e. number of subjects included) and between-run correlation in the group average. The plotted values and error bars represent the mean and standard deviation across all possible combinations. As expected, consistency declines with decreasing group size.

**Figure 4.**
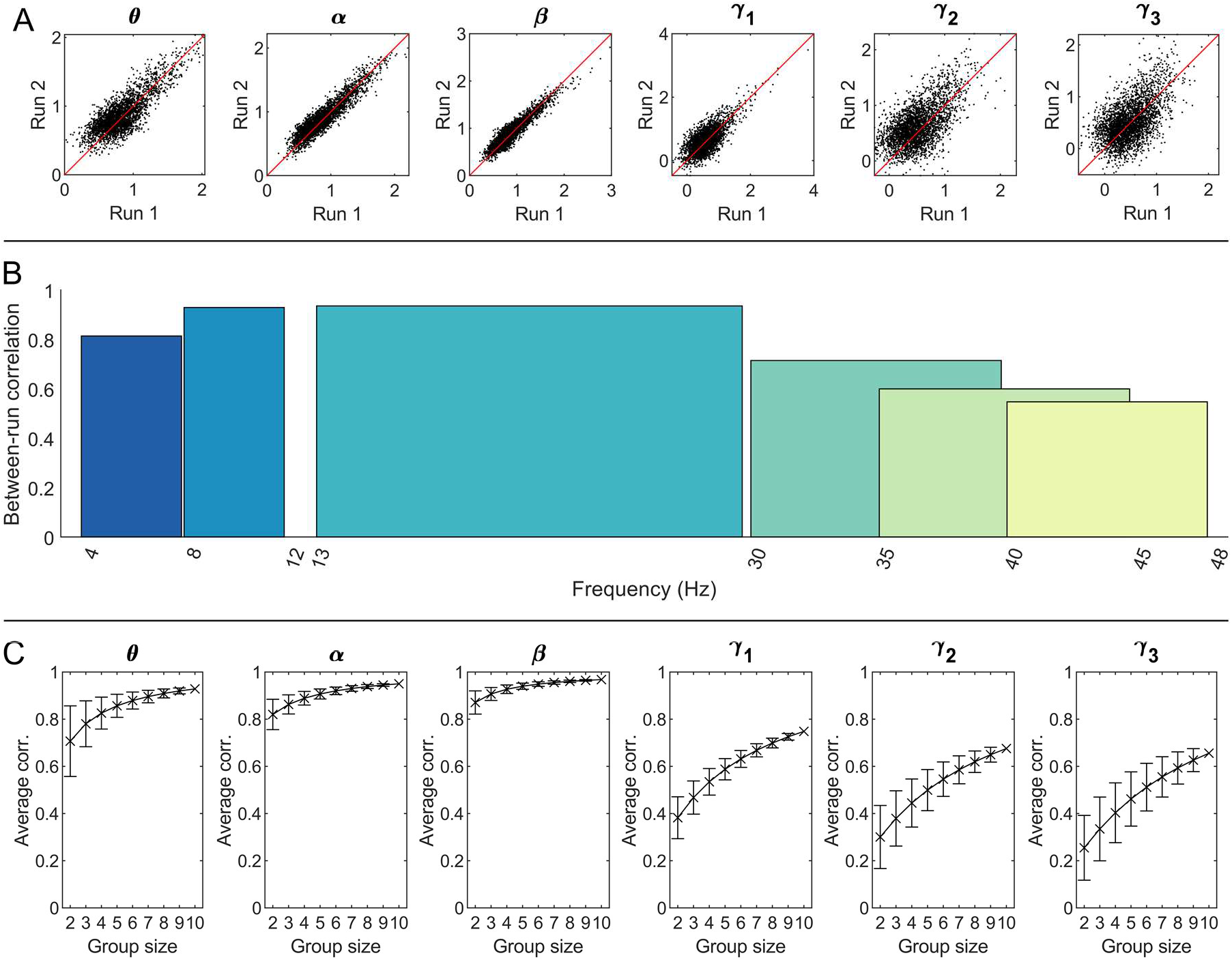
Between-run reliability of the functional connectome. A) Scatter plots of group average connectivity values; run 1 plotted against run 2. Black points represent the mean AEC values for each of the 3003 edges in the group average connectomes for both runs. Lines of equality are indicated in red. B) Bar chart of Pearson correlation coefficients between the across-run average connectomes. Low-frequency connectomes are highly consistent while the gamma sub-bands display more variability between the two experimental runs. C) The effect of sample size on group average between-run correlation. Crosses represent mean correlation values across possible subsamples for each group size; error bars show the standard deviation across subsamples.

Correlation values for our sample size of 10 appear closest to their asymptotic limit in beta, alpha and theta, while group average consistency may be further increased using larger sample sizes for the gamma bands.

### Individual subject comparisons

Figure 5 shows the individual connectomes for all 10 subjects, for the alpha band, for both experimental runs. All matrices are distinctly structured and display a marked difference between subjects. However, the consistency across the two runs within each individual is striking. This qualitative observation is formalised in Figure 6A which shows within- and between-subject correlations between connectome matrices. Recall there are 10 possible within-subject comparisons and 90 between-subject comparisons between runs 1 and 2. In Figure 6A the bars show the mean correlation values whilst the dots show individual values. The difference between within- and between-subject averages is shown in Figure 6B as a function of frequency band. Within-subject correlation (Figure 6C) peaked in the beta band at 0.78 but was high for theta (0.56) and alpha (0.72). In agreement with the group result, it drops for the gamma bands. The within/between-subject difference (Figure 6B) peaked in the alpha band but according to our Monte-Carlo test was significant in the theta, alpha, beta and low gamma bands. In agreement with this, using neural fingerprinting analysis, we were able to correctly identify 7, 10, 8 and 5 individuals in the theta, alpha, beta and low gamma bands respectively, by looking at the highest values of correlation across the group.

**Figure 5:**
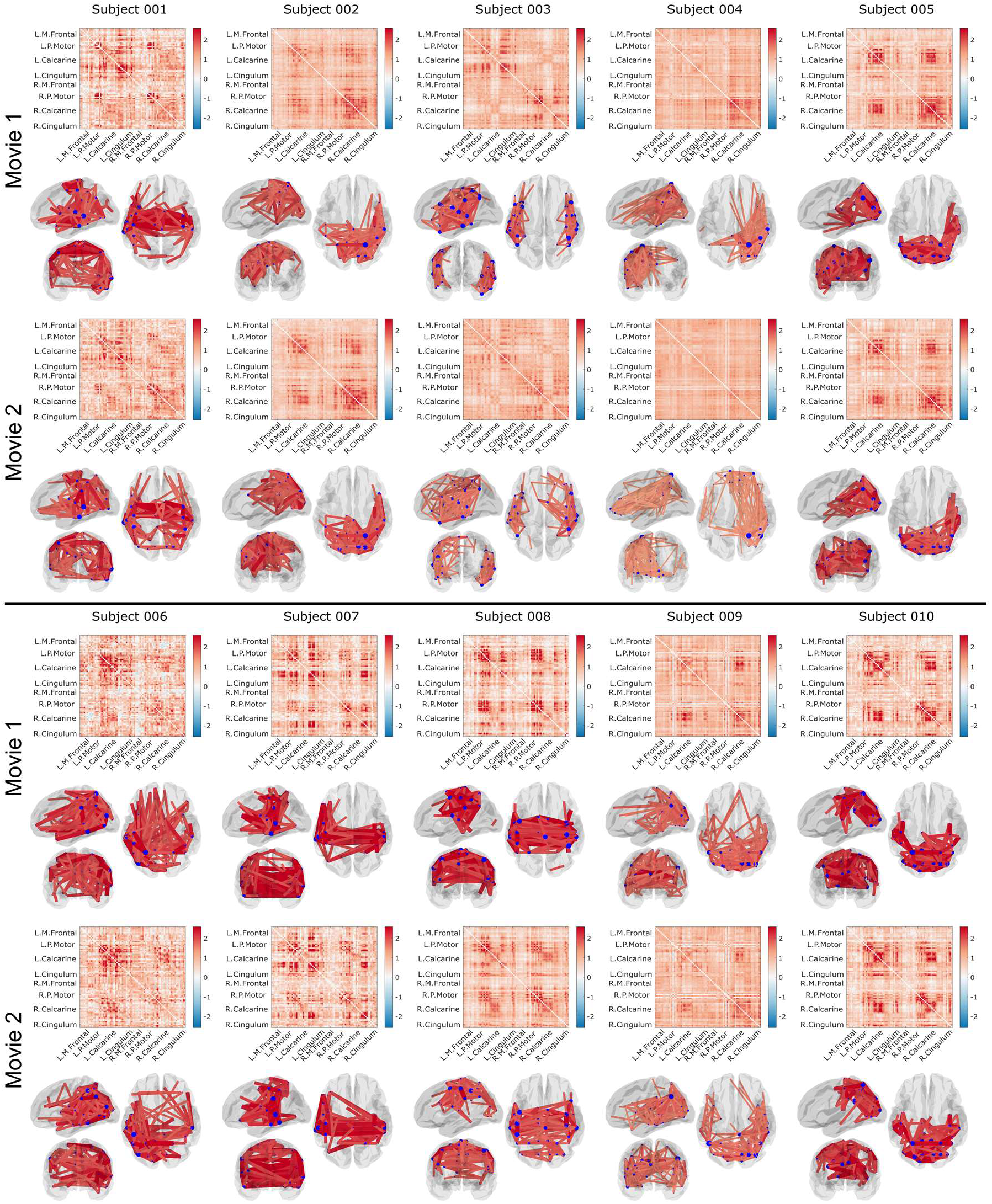
Individual connectivity matrices: Connectomes and corresponding glass brain plots, for all subjects and both experimental runs in the alpha band. Note that whilst variability is high between individuals, results within a single individual are consistent.

**Figure 6:**
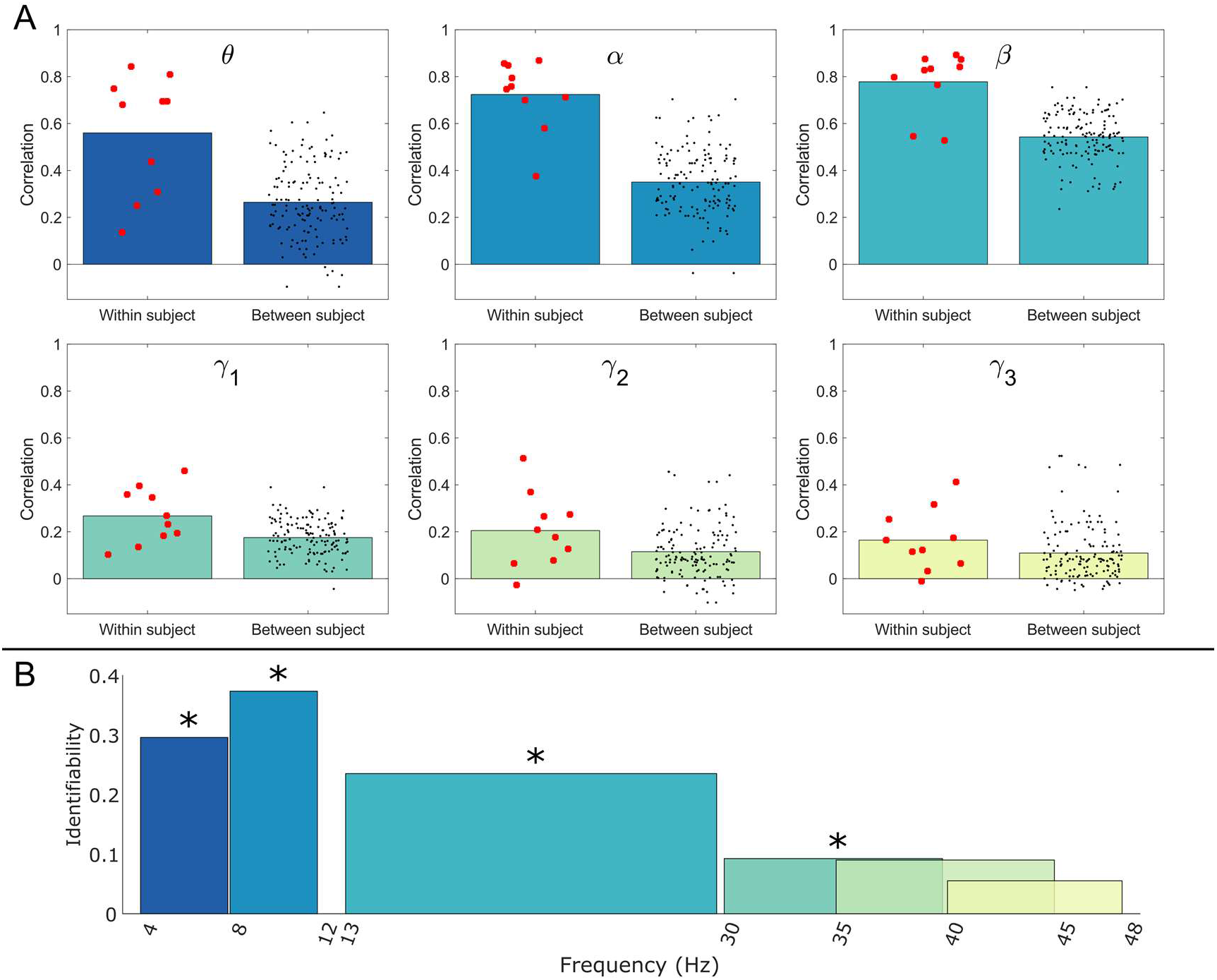
Individual subject comparison across bands. A) Within- and between-subject Pearson correlations of the AEC connectomes for each frequency band. B) Identifiability — the difference between average within- and between-subject correlations indicating the potential for neural fingerprinting across frequency bands. Asterisks indicate statistical significance at the p<0.05 level.P-values were estimated via a permutation test and corrected for multiple comparisons across frequency bands using the Benjamini-Hochberg procedure (Benjamini and Hochberg, 1995).

## DISCUSSION

The ability to characterise functional connectivity reliably is a critical function of any viable MEG system. However, the measurement of connectivity poses a significant challenge as it relies heavily on high-fidelity (unaveraged) MEG data and whole-brain coverage. Here we provide a benchmark for the repeatability of both neural oscillatory activity and connectivity across experimental runs using a 168-channel whole-head OPM-MEG device.

At the group level, the correlation between participant-averaged connectivity matrices for runs 1 and 2 was high in the theta (0.814) alpha (0.929) and beta (0.935) bands. However, this fell to 0.714, 0.599 and 0.547 for *γ*_1_, *γ*_2_ and *γ*_3_ respectively. These values compare well to those previously derived for conventional MEG. Colclough et al. (2016), using groups of ∼30 individuals, demonstrated a between-group correlation of ∼97% in the alpha band using a similar amplitude envelope correlation metric. Whilst values here are marginally lower, this is likely explained by our groups being smaller (10 people). Figure 4C showed that for bands with relatively lower group-level consistency – such as our gamma bands – larger sample sizes can be expected to yield improved consistency. Overall, the high consistency observed in the lower-frequency bands demonstrates that – for group-level measurement – OPM-MEG provides a robust estimate of functional connectivity. The reason for the fall in the gamma band can be seen in Figure 2 when comparing the power spectral density from the brain with that from an empty room noise recording. At frequencies above ∼30 Hz, the “noise” level is around half of the signal amplitude. Above these frequencies, noise begins to dominate and measures become unreliable. This agrees with observations in conventional MEG. For example, a previous study of motor network connectivity (Brookes et al., 2011) showed that connectivity between left and right motor cortices in the resting state was measurable up to ∼40 Hz; similar observations were found in the frontoparietal and default mode networks (Brookes et al., 2011; Hipp et al., 2012).

At the individual level, correlations were lower. Within-subject correlations were 0.56, 0.72 and 0.78 for the theta, alpha and beta bands respectively. In comparison, Colclough et al. (2016b) observed within-subject consistency of ∼58% in the alpha band. This is somewhat lower than the values observed in our study, though it was estimated using shorter segments of data (5 mins rather than 10 mins). Liuzzi et al. (2017) used amplitude envelope correlation applied to conventional MEG, achieving within-subject consistencies of ∼72% in the beta band using 560 s of data and with the head clamped into the MEG helmet to eliminate any motion relative to the (SQUID-based) sensors. In line with expectation, the between-subject consistencies were generally much lower, peaking at 0.54 for the beta band. Once again this is in line with expectations from conventional MEG, with Colclough et al. (2016) showing a between-subject correlation of ∼45% for the alpha band. Based on both the group level and individual observations above, the repeatability of OPM-MEG compares favourably with previously published conventional MEG findings.

The drop in correlation values when undertaking between-subject versus within-subject comparisons is the basis for the technique known as neural fingerprinting. Briefly, successful neural fingerprinting requires that a subject can be correctly identified from a group, based on some feature derived from a previous scan. Here, alpha band connectome matrices enabled successful neural fingerprinting in all ten subjects, with the beta band offering 8 correctly identified individuals and the theta band 7 correctly identified individuals. The gamma band was less successful, and this is also reflected in the fact that the within-subject versus between-subject differences were not significant in y_2_ and y_3_. The topic of neural fingerprinting has gained significant traction in recent years (da Silva Castanheira et al., 2021) with the idea that between-subject variance (which is often treated as noise) contains useful and reproducible information. Indeed, it offers the exciting possibility that, by tracking changes in the neural fingerprint, one might enable early detection of disorders (e.g. dementia). The data presented demonstrate that OPM-MEG is a robust platform from which to launch such studies. At a technical level, there are several limitations of our system which should be addressed. First, the channel count of 168 remains significantly lower than that of conventional MEG systems (which have ∼300 channels). In addition, our triaxial design measures the tangential and radial components of the magnetic field, whereas conventional MEG only measures radially. Whilst the use of triaxial sensors has proven to be an excellent means to reduce the effects of non-brain sources of magnetic field (Brookes et al., 2021; Rea et al., 2022; Tierney et al., 2022), the tangential field components are smaller in amplitude and consequently, in terms of absolute signal, OPM-MEG remains disadvantaged compared to cryogenic instrumentation. It is encouraging that, despite the lower channel count, we achieve approximate parity with conventional MEG in terms of repeatability of connectivity measurement. In addition, one significant advantage of the triaxial design is that three-axis measurement enables complete calibration of the sensor and removal of cross-talk artefacts between close-set sensors. This means that, ostensibly, the construction of high-density whole-head OPM systems should be possible in the near future.

Aside from channel count, one important observation is that, at high frequencies (∼60 Hz), the signal and empty room noise levels begin to converge. Importantly, this does not mean that OPM-MEG cannot assess high-frequency activity; indeed several papers (Hill et al., 2022, 2019; Iivanainen et al., 2020) have shown that OPM-MEG can successfully record gamma band (>50 Hz) oscillations, with similar (or even better) SNR to that observed in conventional MEG (Hill et al., 2020), albeit using tasks specifically designed to induce localised gamma activity It may be somewhat surprising that we did not see strong visual networks in the high frequencies, given that invasive recordings in animals watching natural scenes show significant gamma band activity during naturalistic stimulation (Brunet et al., 2015). However, high-frequency signals may be localised – without strong correlations to distant sources. Whether correlated broadband gamma activity during naturalistic stimulation is detectable using MEG (regardless of sensor noise floor) remains an open question, but enhancing SNR (by decreasing the inherent sensor noise floor) is a core requirement if we are to stand a chance at measuring it. It should be noted that the noise floor of OPMs remains somewhat higher than a SQUID (e.g. for triaxial sensors the noise floor is ∼13 fT/sqrt(Hz), compared to (typically) <5 ft/sqrt(Hz) for SQUIDs). Reducing this is a priority for future OPM implementations.

Finally, in previous conventional MEG studies, data have typically been recorded in the “pure” resting state (i.e. participants are asked to “sit still and do nothing”). In contrast, here, subjects were asked to watch a movie. The addition of this naturalistic stimulus likely drives brain activity which is synchronised across runs; the extent to which this might help to enhance consistency between experimental runs is unknown, though the low between-subject reliability observed (correlation coefficients of 0.35 in the alpha band) would suggest the effect is not large. What is important is that this same movie has been used in previous work to contrast multiple imaging modalities including EEG, fMRI and ECoG (Haufe et al., 2018). Whilst here we intended to measure the consistency of connectome characterisation, it should be the aim of future studies to employ these data and introduce OPM-MEG to a growing comparison of modalities using this same naturalistic stimulus.

## CONCLUSION

OPM-MEG offers significant advantages over conventional MEG, and other non-invasive functional imaging modalities including EEG, fNIRS, and fMRI. However, OPM-MEG is also a new technology. Demonstrating both the viability and repeatability of key metrics is a necessary step in the path to adoption by the neuroimaging community. Here, we aimed to test the robustness of whole-brain connectivity across two separate experimental runs of the same movie-watching paradigm. Results showed that the power spectra of the neural signal, from which connectivity is derived, were consistent across repeats of the experiments, with differences between runs amounting to 4% of the total signal. When assessing connectivity we demonstrated excellent group-level robustness, with high correlations between connectomes in the theta (0.81) alpha (0.93) and beta (0.94) frequency ranges. At the individual subject level, we found marked differences between individuals, but high within-subject robustness (correlations of 0.56 ± 0.25, 0.72 ± 0.15 and 0.78 ± 0.13 in theta, alpha and beta respectively). These results compare well to equivalent findings using conventional MEG; they show that OPM-MEG is a viable way to characterise whole-brain connectivity and add significant weight to the argument that OPMs can overtake cryogenic sensors as the fundamental building block of MEG systems.

## ACKNOWLEDGEMENTS

This work was supported by an Engineering and Physical Sciences Research Council (EPSRC) Healthcare Impact Partnership Grant (EP/V047264/1) and an Innovate UK germinator award (Grant number 1003346). We acknowledge support from the UK Quantum Technology Hub in Sensing and Timing, funded by EPSRC (EP/T001046/1). Sensor development was made possible by funding from the National Institutes of Health (R44MH110288). S.M. is funded by Deutsche Forschungsgemeinschaft (DFG), project 437219953. We also express thanks to Cerca Magnetics Ltd., Metamorphic AM and Added Scientific Ltd. for the useful and productive discussions that led to the design of the lightweight helmet used for paediatric measurements.

## DATA AND CODE AVAILABILITY

All data used to produce the results presented here will be made available upon acceptance of the manuscript at https://doi.org/10.5281/zenodo.7477061. The MATLAB software used for data analysis will be available at https://github.com/LukasRier/Rier2022_OPM_connectome_test-retest/.

